# Characterization of genomic variants associated with resistance to bedaquiline and delamanid in naïve *Mycobacterium tuberculosis* clinical strains

**DOI:** 10.1101/2020.05.27.120451

**Authors:** S Battaglia, A Spitaleri, AM Cabibbe, CJ Meehan, C Utpatel, N Ismail, S Tahseen, A Skrahina, N Alikhanova, Kamal SM Mostofa, A Barbova, S Niemann, R Groenheit, AS Dean, M Zignol, L Rigouts, DM Cirillo

**Affiliations:** Emerging Bacterial Pathogens Unit, Division of Immunology, Transplantation and Infectious Diseases, IRCCS San Raffaele Scientific Institute, Milan, Italy; Center for Omics Sciences, IRCCS San Raffaele Scientific Institute, Milan, Italy; School of Chemistry and Biosciences, University of Bradford, Bradford, UK; Unit of Mycobacteriology, Department of Biomedical Sciences, Institute of Tropical Medicine, Antwerp, Belgium; Molecular and Experimental Mycobacteriology, Research Center Borstel, Borstel, Germany; Centre for Tuberculosis, National Institute for Communicable Diseases, National Health Laboratory Services, Johannesburg, South Africa; National TB Reference Laboratory, National Tuberculosis Control Program, Islamabad, Pakistan; Republic Research and Practical Centre for Pulmonology and Tuberculosis, Minsk, Belarus; Scientific Research Institute of Lung Diseases, Ministry of Health, Baku, Azerbaijan; National TB Reference Laboratory, National Institute of Diseases of the Chest and Hospital, Dhaka, Bangladesh; Central Reference Laboratory on Tuberculosis Microbiological Diagnostics, Ministry of Health, Kiev, Ukraine; German Center for Infection Research (DZIF), Partner Site Hamburg-Lübeck-Borstel-Riems, Germany; Unit for Laboratory Surveillance of Bacterial Pathogens, Public Health Agency of Sweden, Solna, Sweden; Global TB Programme, World Health Organization, Geneva, Switzerland; Department Biomedical Sciences, Antwerp University, Antwerp, Belgium

## Abstract

The role of genetic mutations in genes associated to phenotypic resistance to bedaquiline (BDQ) and delamanid (DLM) in *Mycobacterium tuberculosis* complex (MTBc) strains is poorly understood. However, a clear understanding of the role of each genetic variant is crucial to guide the development of molecular-based drug susceptibility testing (DST). In this work, we analysed all mutations in candidate genomic regions associated with BDQ- and DLM-resistant phenotypes using a whole genome sequencing (WGS) dataset from a collection of 4795 MTBc clinical isolates from six high-burden countries of tuberculosis (TB). From WGS analysis, we identified 61 and 158 unique mutations in genomic regions potentially involved in BDQ- and DLM-resistant phenotypes, respectively. Importantly, all strains were isolated from patients who likely have never been exposed to the medicines. In order to characterize the role of mutations, we performed an energetic *in silico* analysis to evaluate their effect in the available protein structures Ddn (DLM), Fgd1 (DLM) and Rv0678 (BDQ), and minimum inhibitory concentration (MIC) assays on a subset of MTBc strains carrying mutations to assess their phenotypic effect. The combination of structural protein information and phenotypic data allowed for cataloging the mutations clearly associated with resistance to BDQ (*n=* 4) and DLM (*n=* 35), as well as about a hundred genetic variants without any correlation with resistance. Importantly, these results show that both BDQ and DLM resistance-related mutations are diverse and distributed across the entire region of each gene target, which is of critical importance to the development of comprehensive molecular diagnostic tools.

**Importance:** Phenotypic drug susceptibility tests (DST) are too slow to provide an early indication of drug susceptibility status at the time of treatment initiation and very demanding in terms of specimens handling and biosafety. The development of molecular assays to detect resistance to bedaquiline (BDQ) and delamanid (DLM) requires accurate categorization of genetic variants according to their association with phenotypic resistance. We have evaluated a large multi-country set of clinical isolates to identify mutations associated with increased minimum inhibitory concentrations (MICs) and used an *in silico* protein structure analysis to further unravel the potential role of these mutations in drug resistance mechanisms. The results of this study are an important source of information for the development of molecular diagnostic tests to improve the provision of appropriate treatment and care to TB patients.

## 1. Introduction

The management of drug resistant tuberculosis (DR-TB) caused by *Mycobacterium tuberculosis* complex (MTBc) strains poses a serious public health challenge worldwide. The World Health Organization (WHO) new guidelines recommend the use of bedaquiline (BDQ) for all TB cases of rifampicin resistance (RR-TB), multi drug resistant (MDR) TB (MTBc strains resistant at least to isoniazid and rifampicin) and extensively drug-resistant (XDR) TB (MDR strains resistant to fluoroquinolones and second-line injectables drugs) (**1**). Based on WHO priority grouping of medicines, Delamanid (DLM) compound is recommended when an effective regimen cannot be established by group A agents, containing BDQ, fluoroquinolones (FQs), linezolid (LZD) and B agents group, composed by clofazimine (CLF) and cycloserine (CS) (**2**). Previous studies have shown that MDR/XDR-TB patients treated with BDQ and DLM rapidly develop resistance due to fixed mutations in candidate genes which often appear as previously undescribed novel variants (**3, 4, 5, 6, 7**). Additionally, resistance to BDQ can arise in naïve populations of MTBc strains as a consequence of a clofazimine-containing regimen, or by random mutations affecting the drug targets (**8, 9, 10, 11, 12, 13, 14**). DLM resistance not associated to exposure has been reported as well (**15, 16, 17, 18, 19**). Moreover, as a member of the nitroimidazooxazines, DLM shares the same resistance mechanisms of pretomanid (PA-824) compound (**20**).

Therefore, knowledge of the BDQ and DLM susceptibility status of clinical MTBc isolates before therapy has started and the early detection of emerging resistance in failing regimens are needed to ensure an effective treatment of DR-TB.

Here, whole genome sequencing (WGS) based approaches, that are rapidly expanding from basic research into routine diagnostic laboratories, provide the advantage of interrogate virtually all resistance targets in a given clinical MTBc strain. However, the routing diagnostic application of WGS requires a much better understanding of the correlation between genotypic and phenotypic, particularly for the new drugs (**21, 22**).

Currently, the molecular mechanisms leading to resistance to BDQ and DLM are not well described, a fact that jeopardises the design of a reliable molecular approach to detect resistance.

Mutations in *atpE* (*Rv1305*), which encodes for the C-subunit of ATP synthase, have been associated with phenotypic resistance to BDQ, which is known to directly inhibit the ATP synthase (on target mechanism) (**23**). In addition, the mutations in the off-target *Rv0687* gene result in increased minimum inhibitory concentrations (MICs) for BDQ by up-regulation of the MmpL5/MmpS5 pump gene expression, concurrently leading to a cross-resistance to clofazimine (CLF) (**10, 11, 24**). Furthermore, it has been demonstrated that mutations in *pepQ* (*Rv2535c*) may confer low-level resistance to both CLF and BDQ in clinical isolates (**9, 25**).

DLM impairs the biosynthesis of mycolic acids and requires activation by the F420-dependent nitroreductase encoded by the *ddn* gene (on target mechanism). The F420 cofactor is synthetized by enzymes encoded by the *fbiA, fbiB, fbiC, fgd1* genes, all of which are involved in DLM off-target mechanisms. Polymorphisms in these genes were shown to be involved in phenotypic DLM resistance (**26, 27**).

To better describe BDQ and DLM resistance mechanisms, we investigated the genomic regions involved in resistance to BDQ and DLM from 4795 MTBc isolates collected within a multi-country drug-resistance surveillance study and to identify variants potentially involved in resistance development (**28**). The mutations found were correlated with strain lineage, DR-profile and country of origin. To combine the genomic data with phenotypes, we performed BDQ and DLM MICs for a subset of these isolates. Finally, for the available 3D protein structures (Ddn, Fgd1 and Rv0678) we performed an *in silico* structural and energetic analysis in order to characterize and quantify the mutation effect on protein function. This combined information enabled us to provide a first robust catalogue of BDQ and DLM resistance mutations as basis of the establishment of WGS resistance prediction algorithms for these drugs.

## 2. Methods

### 2.1 Study design

A total of 4795 genome sequences retrieved from the Sequence Read Archive of the National Center for Biotechnology Information as recalibrated BAM files (accession number SRP128089) were considered and investigated in this study. The corresponding MTBc isolates originate from a unique population-based surveillance study across six countries with a high burden of TB or MDR-TB, according to WHO’s high burden country list for the period of 2016-2020: Azerbaijan (*n* = 751), Belarus (*n* = 197), Bangladesh (*n* = 935), Pakistan (*n* = 194), South Africa (*n* = 1578), Ukraine (*n* = 1140) (**28**). For our purposes, all sequenced isolates harbouring at least one single nucleotide polymorphism (SNP) or insertion/deletion (indel) in at least one of the candidate genomic regions for DLM and/or BDQ resistance were considered for the analysis, excluding synonymous mutations and previously characterized lineage-associated SNPs for which the absence of correlation with the phenotypic DLM resistance was demonstrated (**15, 16**). For genetic variants detected in more than one isolate we decided to replicate results selecting two isolates from different countries, whenever possible. The flowchart for sample selection, the number of isolate tested and phenotypic drug-susceptibility testing (DST) of selected isolates are reported in Figure S1 of supplementary materials.

### 2.2 Whole Genome Sequencing analysis

WGS data were generated by both Illumina technology (Illumina, San Diego, CA, USA) and Ion Torrent technology (ThermoFisher Scientific) as previously described (**28**). Sequencing data were analysed using the MTBseq pipeline (**29**) to identify all variants in the genomes and MTBc lineage. The analysis was performed on the mapped MTBc reads by setting a quality threshold of at least a mean coverage of 20x and an unambiguous base call threshold of ≥70%. A mutation was called only if SNPs and/or indel variants were detected by at least eight reads (both forward and reverse reads) with a minimum phred quality score of 20, and by considering a mutation frequency of ≥75%. The regions of the MTBc genome reference H37Rv NC_000962.3 (**30**) considered in the study are reported in Table S1 of supplementary materials. The WGS analysis results and distribution of mutations among lineages and countries of isolation are reported in the supplementary excel Dataset S1.

Cluster analysis was performed on the distance matrix generated by the MTBseq pipeline using *in-house* python scripts (https://github.com/aspitaleri/python). The distance matrix was analysed using a hierarchical linkage clustering method with a 12 SNPs cut-off (**31**).

### 2.3 Minimum Inhibitory Concentration assay

The selected MTBc isolates for genetic variants were sub-cultured on Löwenstein-Jensen medium and subsequently subjected to MIC testing against BDQ and/or DLM by the resazurin colorimetric microtiter plate assay (REMA) as previously described (**16, 32, 33**). Delamanid powder was obtained from Otsuka Pharmaceutical (Tokyo, Japan) and pure bedaquiline powder was obtained from Janssen Pharmaceutical (Beerse, Belgium). A DLM concentration range of 0.004-4 µg/ml and a BDQ concentration range of 0.004-2 µg/ml were used, considering the proposed cut-off values of 0.12 µg/ml and 0.06 for BDQ and DLM, respectively (**34**). Based on MIC results, the isolates were categorized as susceptible (S; MIC ≤ cut-off), low resistant level (I; MIC 1 dilution > cut-off) or resistant (R; MIC more than 1 dilution > cut-off). All MIC values reported in this work correspond to the MIC100 value that considers any change in colour to purple/pink as indicating the presence of viable bacilli (**33**). For each batch of isolates tested, the H37Rv *M. tuberculos*is reference strain (*M. tuberculosis* H37Rv ATCC 27294) was included as a control, and test isolate results of that batch were accepted only if the H37Rv MIC value was within the expected range of ≤0.004-0.03 µg/ml for DLM and ≤0.008-0.03 µg/ml for BDQ. Further details of REMA protocols are reported in supplementary materials Text S1 word file.

### 2.4 Mutation structural analysis

We carried out an energetic analysis on the available crystal structures of proteins Ddn (PDB 3R5R), Fgd1 (PDB 3B4Y) and Rv0678 (PDB 4NB5) (**35, 36, 37**) using Eris (**38**) and MAESTRO (**39**) programs. We exploited two end-point methods to evaluate the change of protein stability upon mutations, namely Eris and MAESTRO, which calculate the folding free energy in two different manners. For this structural analysis stop codons, frameshifts, and SNPs affecting the promoter region were not included. The stability change, ΔΔG, is computed as the difference between the average stabilities of mutant and wild type protein structures. Both *in silico* approaches were used as a qualitative cross-validation to evaluate the protein mutation effects, considering 0.34 Kcal/mol and 5 Kcal/mol as thresholds for MAESTRO and Eris, respectively.

In addition to folding stability, we calculated the effect of mutations on the complex stabilities Ddn-F420-H_2_ Fgd1-F420-H_2_ using DSX pair potentials knowledge-based scoring function (**40**). In case of Rv0678 protein we also performed an energetic analysis to quantify the effect of the mutations on the homodimer protein-protein stability. For this purpose, we carried out a molecular mechanics energies combined with the Generalized Born and surface area continuum solvation (MM/GBSA). The calculation was performed using MMPBSA.py program within Amber14 suite using ff14 force field and the GB^OBC1^ implicit solvent model (**41**). All obtained *in silico* results are reported in the supplementary material Dataset S2. Further details of *in silico* analyses are reported in supplementary materials Text S1 word file. Primary protein sequences alignment for the frameshift analysis was performed using ClustalX algorithm (**42**). The visualization of the mapped mutations on the protein structures are created with PyMOL v2.0 (**43**).

## 3. Results

A total of 4795 WGS from MTBc clinical strains were analysed by considering the candidate genomic regions for BDQ resistance (*atpE, Rv0678, pepQ*) and for DLM resistance (*ddn, fgd1, fbiA, fbiB, fbiC*). This collection included 731 (17%) MDR and 79 (2%) XDR MTBc strains (Fig. S1). Based on WGS results, we identified a total of 106 and 643 isolates harboring relevant genomic variants potentially involved in BDQ and DLM resistance, respectively. We tested a subset of isolates carrying mutations for phenotypic DST for BDQ (*n=* 51) and for DLM (*n=* 124) representing the 43 and 104 BDQ- and DLM-related variants, respectively (Fig. S1). All genomic variants detected by WGS analysis in candidate genes for BDQ and DLM with the corresponding information of MTBc strain lineage, drug resistance profile, country of isolation, mutation frequency and MIC results for tested MTBc isolates are reported in Dataset S1 of supplementary material.

### 3.1. Analysis of mutations for BDQ resistance

The WGS analysis revealed 61 unique mutations in the considered genomic regions associated with BDQ resistance (Fig. S1). The mutation analysis distribution revealed 27 unique mutations in *Rv0678* (including 7 mutations in the promoter region, 16 nonsynonymous mutations, and 4 indels causing frameshift mutations), 32 unique mutations in the *pepQ* gene (including 2 upstream mutations, 28 nonsynonymous mutations and 2 frameshift mutations), and 2 upstream mutations in the *atpE* gene, while no mutations were found in the *atpE* encoding region (Dataset S1).

Phenotypic testing revealed that four different *Rv0678* mutations had MIC values above the cut-off of 0.12 µg/ml: two frameshift (fs) mutations in *Rv0678*, Gly6fs (del_16-17 gg) and the double mutant Gln9fs-Thr92fs (ins_27 c, ins_274 a) associated with an MIC of 0.5 µg/ml, and two *Rv0678* nonsynonymous mutations Arg96Trp and Met111Lys yielding a low level of resistance to BDQ of 0.25 µg/ml (Table 1). The two frameshift mutations associated with BDQ-resistant phenotypes were observed in one MDR isolate and another MDR isolate with concurrent resistance to FQs (corresponding to one new and one previously treated TB case, respectively). Two nonsynonymous mutations associated to low level resistance were found in two pan-susceptible (full-S) MTBc strains (Table 1). The other 39 mutations were detected in isolates susceptible to BDQ with MIC values ≤0.12 µg/ml, including two isolates harbouring frameshifts in Rv0678: Ile16fs (ins_ 46 tcatggaattcg) and Ala153fs (ins_457 c) showing an MIC of 0.06 µg/ml. (Dataset S1). The protein amino acid sequences obtained from these two frameshifts mutations were aligned to the Rv0678 wild-type sequence, highlighting that both wild-type and mutated proteins contain the two well-conserved and important regions: the amino acid (aa) positions from 34 to 99 (DNA-binding domain) and positions from 16 to 32 and from 101 to 160 (dimerization domains) (Fig. 1). In the case of *Rv0678* Ile16fs, the insertion of 12 nucleotides caused the addition of 4 aa from position 16 of the Rv0678 protein without disrupting the frame of the whole enzyme, while the Ala153fs caused a change to the last 13 aa of the C-terminal of the protein (Fig 1). This suggests that these frameshifts do not affect protein stability and function resulting in the BDQ-susceptible phenotype.

**Table 1.**
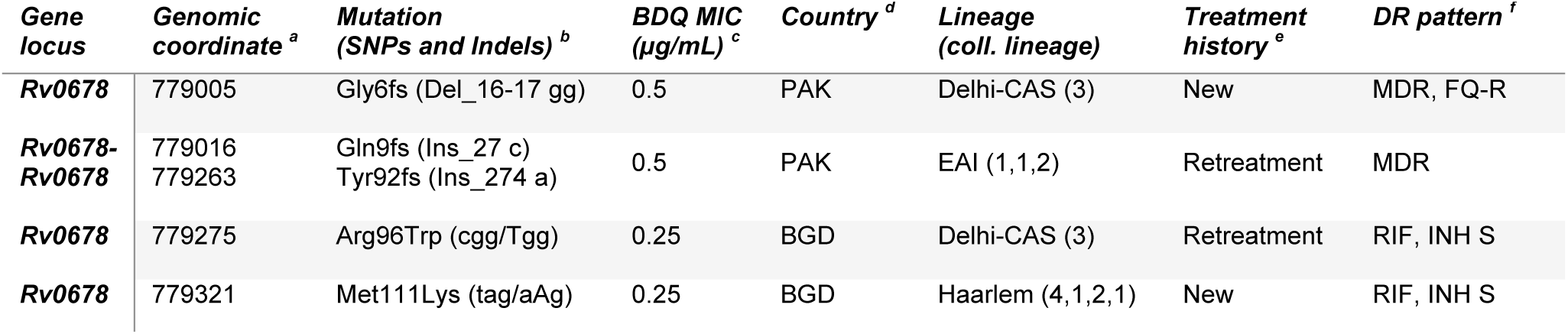
List of *Rv0678* mutations detected in MTBc strains resistant to BDQ. In the table are reported all the information for each BDQ-resistant related mutations (MIC ≥ 0.25 µg/ml) and for the MTBc isolate tested for BDQ susceptibility. ^a^ Genomic position in reference H37Rv NC_000962.3 strain; ^b^ Amino acid (aa) change and nucleotide (nt) change; ^c^ Minimum Inhibitory concentration (MIC) value in µg/ml. ^d^ Country of MTBc isolate origin: Pakistan (PAK), Bangladesh (BGD). ^e^ TB patient treatment history: new TB case “New”, or patient with previous TB history and treatment “Retreatment”. ^f^ Drug resistance pattern of the isolates: MDR (multi drug resistant strain), FQs (fluoroquinolones), INH (Isoniazid), RIF (Rifampicin), R/S (resistant/susceptible).

| *Gene locus* | *Genomic coordinate<sup>a</sup>* | *Mutation (SNPs and Indels)<sup>b</sup>* | *BDQ MIC (µg/mL)<sup>c</sup>* | *Country<sup>d</sup>* | *Lineage (coll. lineage)* | *Treatment history<sup>e</sup>* | *DR pattern<sup>f</sup>* |
| --- | --- | --- | --- | --- | --- | --- | --- |
| *Rv0678* | 779005 | Gly6fs (Del_16-17 gg) | 0.5 | PAK | Delhi-CAS (3) | New | MDR, FQ-R |
| *Rv0678-Rv0678* | 779016<br>779263 | Gln9fs (Ins_27 c)<br>Tyr92fs (Ins_274 a) | 0.5 | PAK | EAI (1,1,2) | Retreatment | MDR |
| *Rv0678* | 779275 | Arg96Trp (cgg/Tgg) | 0.25 | BGD | Delhi-CAS (3) | Retreatment | RIF, INH S |
| *Rv0678* | 779321 | Met111Lys (tag/aAg) | 0.25 | BGD | Haarlem (4,1,2,1) | New | RIF, INH S |

**Fig. 1.**
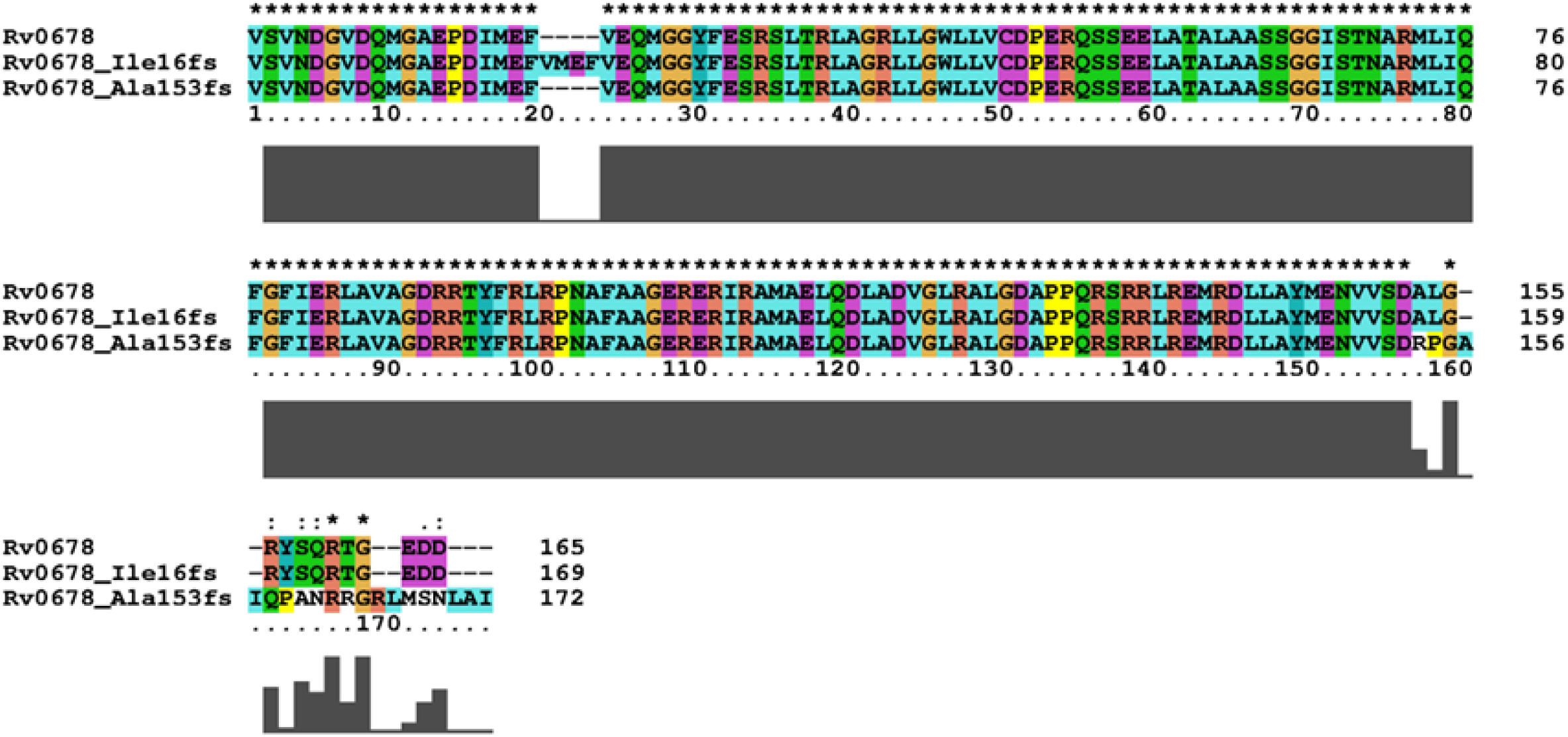
Amino acid sequence alignment of BDQ-susceptible frameshift mutations in *Rv0678*. The amino acid (aa) sequence of the translated proteins with Ile16fs and Ala153fs frameshift mutations (associated to BDQ-susceptible MIC) were aligned with Rv0678 wild type aa sequence using ClustalX. In both cases the two insertions mutations cause the addition of new aa residues without altering the frame of Rv0678 protein and functional residues of the protein.

A structural analysis of the effect of mutations in Rv0678 resulting in amino acid changes was performed as previously described by the Eris and MAESTRO computational approaches (Dataset S2). The Rv0678 folding stability calculation by ERIS software showed that both Arg96Trp and Met111Lys mutations associated with the BDQ-resistant phenotype altered Rv0678 protein folding/stability (ΔΔG kcal/mol >5). These two mutations are localized in the dsDNA-binding and dimerization domain regions of Rv0678, respectively (Fig. 2). Interestingly, the Met111Val mutation had a milder effect on the protein stability than Met111Lys which is reflected in the low BDQ MIC value. Both Eris and MAESTRO analysis showed that the other mutations in Rv0678 have a lower estimated effect on protein stability which is in accordance with lower MIC values for these clinical strains (Dataset S2). As these approaches do not consider the effect of the mutations on dimerization protein function, we calculated the protein-protein binding free energy under the MM/GBSA approximation for the mutations which localize in the Rv0678 dimerization domain (Fig. 2). The results showed that five mutations (Met111Lys, Leu117Arg, Val120Met, Asp141His and Met146Arg) have a significant increase of ΔΔG kcal/mol in the protein-protein homodimer binding free energies, indicating that they can affect the dimerization process, destabilizing the overall homodimer stability (Dataset 2). This data suggest that these mutations could be directly involved in the slight increase in BDQ MIC for these strains, all with a BDQ MIC of 0.12 µg/ml except for the MTBc strain harboring the *Rv067*8 Met146Arg variant with a BDQ MIC of 0.06 µg/ml. (Fig. 2).

**Fig. 2.**
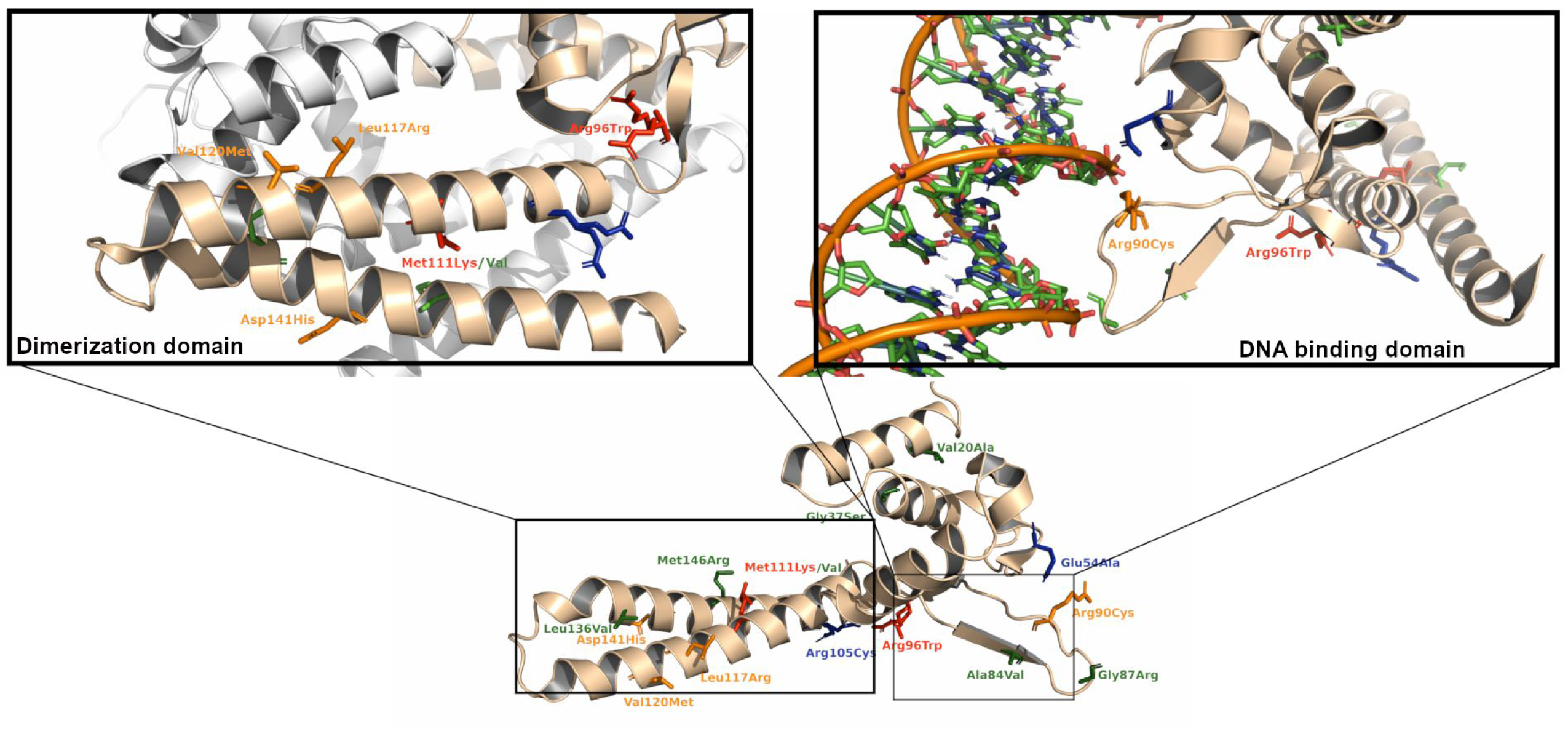
Cartoon representation of Rv0678 protein structures with mutations associated to BDQ-resistant and to BDQ-susceptible phenotypes. Carton representation of the monomer present in the X-Ray unit cell (4NB5). Highlighted in sticks are reported the mutations associated to BDQ resistance (red) and susceptible (green) from BDQ MIC assay. *In silico* predicted mutations which could alter dimerization or DNA binding function, and predicted associated to BDQ-susceptible phenotype are shown in orange and in blue respectively. Cartoon zoomed representations of Rv0678 dimerization domain and DNA binding domain are reported.

An analysis of correlation between observed mutations in *Rv0678, atpE* and *pepQ* regions with lineage and country of origin, revealed that the majority of mutations (*n=* 46; 75.4%) occurred only once (Fig. 3). Only four mutations, all in *Rv0678* gene, were detected in more than 5 isolates, all of them showing a BDQ-susceptible phenotype: the a-4t mutation in the promoter region of *Rv0678* was found in 12 Beijing (2,2,1) isolates from Azerbaijan, the Val3Ile mutation in 8 LAM (4,3,4,2) isolates from Bangladesh, Asn4Thr in 6 Delhi-CAS (3) isolates from Pakistan and Gly87Arg in 8 EAI (1,1,2) isolates from Bangladesh and Pakistan (Fig. 3).

**Fig. 3.**
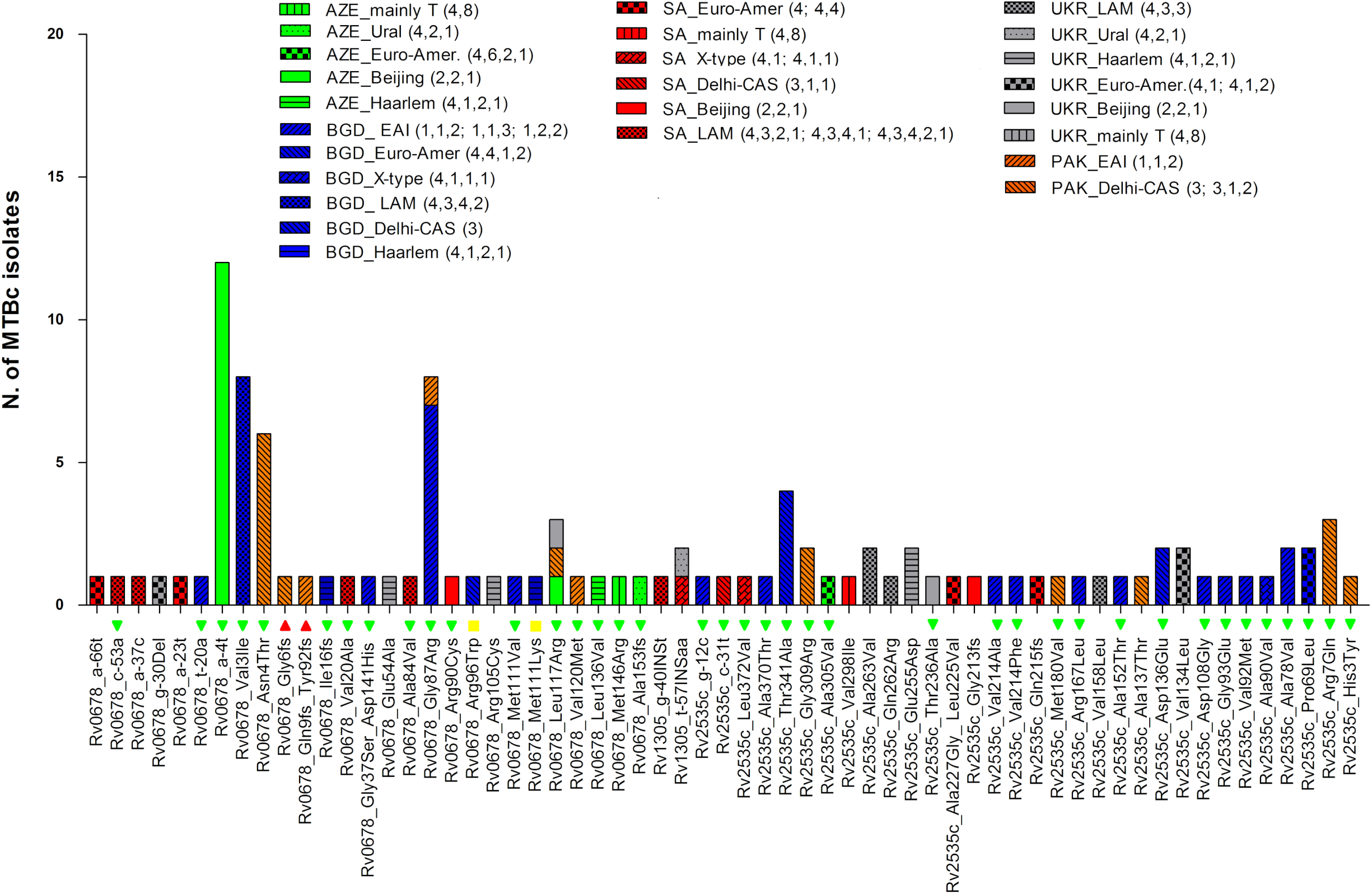
Lineage and country distributions of MTBc strains with variants in *Rv0678, atpE* and *pepQ* genomic regions. The graph reports all mutations found in *Rv0678, atpE* and *pepQ* genomic regions showing their distribution among lineages and country of isolation. The histograms refer to the number of strains in which mutations were observed (Y axis). The colors of the histograms represent the different countries of isolation while the patterns inside each bar represent the different lineages. On the X axis, the results of the MIC test for available MTBc strains are also reported: red triangle are BDQ-resistant strains (MIC > 0.25 µg/ml); yellow box, are BDQ low resistance level strains, (MIC = 0.25 µg/ml); green triangles are BDQ-susceptible strains (MIC ≤ 0.12 µg/ml).

Looking at the BDQ-resistance associated variants, the two frameshift mutations associated with high level of BDQ MICs were both observed in isolates from Pakistan and belonged to EAI (1,1,2) and Delhi-CAS (3) lineages, while the two *Rv0678* mutations associated with low level of BDQ resistance, Arg96Trp and Met111Lys, were both observed in two isolates from Bangladesh belonging to Delhi-CAS (3) and Haarlem (4,2,1,2) lineages (Fig. 3).

### 3.2. Analysis of mutations for DLM resistance

The 643 MTBc isolates identified by WGS as harbouring at least one mutation in one of the candidate genes for DLM resistance represented 164 unique DLM-related mutations. The WGS analysis revealed 30 unique mutations in *ddn* (including 3 nonsense and 2 frameshift mutations), 25 unique mutations in *fbiA* (including 1 nonsense mutation), 23 unique mutations in *fbiB*, 65 unique mutations in *fbiC* (including 2 frameshift mutations) and 24 unique mutations in *fgd1* gene (Dataset S1).

Considering all unique mutations, twenty (12.2%) were combinations of two or three variants in more than one candidate gene. Phenotypic results revealed that out of the 124 isolates tested for DLM, 26 (21%) were resistant to DLM, 13 (10.5%) showed a low level of resistance (MIC = 0.12 µg/ml), while 85 (68.5%) were DLM susceptible (Dataset S1). The DLM-resistant isolates spanned 32 different mutations (Table 2). Considering the phenotypic drug resistance profiles of the DLM-resistant isolates, only six were MDR-TB strains, five of which were retreatment TB cases and one was a new TB case. The analysis of mutation types associated with DLM-resistant revealed 3 nonsense mutations leading to truncated proteins, 3 frameshift mutations and 26 nonsynonymous mutations leading to a single amino acid change (Table 2).

**Table 2.** List of *ddn, fgd1, fbiA, fbiB* and *fbiC* mutations detected in MTBc strains resistant to DLM. In the table are reported all the information for each DLM-resistant related mutations (MIC ≥ 0.12) and for the MTBc isolates tested for DLM susceptibility. ^a^ Genomic position in reference H37Rv NC_000962.3 strain. ^b^ Amino acid change and nucleotide (nt) change; ^c^ minimum Inhibitory concentration (MIC) value in µg/ml. ^d^ Country of MTBc isolate origin: Pakistan (PAK), Bangladesh (BGD), Ukraine (UKR), Azerbaijan (AZE), South Africa (SA). ^e^ Patient treatment history; new TB case “New”, or patient with previous TB history and treatment “Retreatment”. ^f^ Drug resistance pattern of the isolates: MDR (multi drug resistant strain), FQs (fluoroquinolones), INH (Isoniazid), RIF (Rifampicin), R/S (resistant/susceptible).

| *Gene locus* | *Genomic coordinate<sup>a</sup>* | *Mutation (SNPs and Indels)<sup>b</sup>* | *DLM MIC (µg/mL)<sup>c</sup>* | *Country<sup>d</sup>* | *Lineage (coll. Lineage)* | *Treatment history<sup>e</sup>* | *DR pattern<sup>f</sup>* |
| --- | --- | --- | --- | --- | --- | --- | --- |
| *ddn* | 3986846 | Met1fs (Del_2 t) | 2 | BGD | Eur-Amer.(4,1,2) | New | RIF, INH S |
| *ddn* | 3986848 | Pro2Gln (ccg/cAg) | 0.25 | AZE | mainly-T (4,8) | New | RIF, INH S |
| *ddn* | 3986885 | Ser14fs (Del_41 g) | > 4 | BGD | Beijing (2,2,1) | Retreatment | MDR |
| *ddn-fbiA* | 3986923<br>3640445 | Trp27Stop (tgg/tAg)<br>prom (g-18a) | > 4 | BGD | EAI (1,1,3) | Retreatment | MDR |
| *ddn* | 3986932 | Arg30His (cgc/cAc) | 1 | BGD | EAI (1,1,3) | Retreatment | RIF, INH S |
| *ddn* | 3986944 | Gly34Glu (ggg/gAg) | 0.12 | AZE | LAM (4,3,3) | New | RIF, INH S |
| *ddn* | 3987015 | Gln58Stop (cag/Tag) | > 4 | UKR | Beijing (2,2,1) | Retreatment | MDR, FQ-R |
| *ddn-fbiA* | 3987025<br>3641164 | Val61Gly (gtc/gGc)<br>Ile208Val (atc/Gtc) | 0.12 | SA | Eur-Amer.(4,1,2) | New | RIF, INH S |
| *ddn-fbiA* | 3987115<br>3640714 | Asn91Thr (aac/aCc)<br>Val58Ile (gtc/Atc) | 0.25 | AZE | mainly-T (4,8) | New | RIF, INH S |
| *ddn* | 3987262 | Thr140Ile (acc/aTc) | 0.5 | BGD | S-type (4,4,1,1) | New | RIF, INH S |
| *fgd1* | 491723 | Gly314Glu (gga/gAa) | 0.12 | BGD | Beijing (2,2,1) | New | RIF, INH S |
| *fbiA* | 3640546 | Lys2Glu (aag/Gag) | 0.12 | AZE | Beijing (2,2,1) | Retreatment | RIF, INH S |
| *fbiA* | 3641002 | Val154Ile (gta/Ata) | 0.12 | BGD | Beijing (2,2,1) | Retreatment | MDR, FQ-R |
| *fbiA-fbiB* | 3641018<br>3642877 | Pro159Gln (ccg/cAg)<br>Lys448Arg (aag/aGg) | 0.12 | BGD | Delhi-CAS (3) | Retreatment | RIF, INH S |
| *fbiA* | 3641164 | Ile208Val (atc/Gtc) | 0.25 | BGD | Eur-Amer.(4,1,2) | New | RIF, INH S |
| *fbiA* | 3641167 | Ile209Val (atc/Gtc) | 0.5 | BGD | Eur-Amer.(4,5) | New | RIF, INH S |
| *fbiA* | 3641403 | Cys287Stop (tgc/tgA) | 4 | PAK | EAI (1,1,2) | New | RIF, INH S |
| *fbiA* | 3641453 | Arg304Gln (cgg/cAg) | 0.25 | PAK | Delhi-CAS (3) | New | MDR |
| *fbiB* | 3642195 | Gly221Ser (ggc/Agc) | 0.12 | BGD | Beijing (2,2,2) | New | FQ-R |
| *fbiB* | 3642204 | Asp224Asn (gac/Aac) | 0.5 | BGD | Delhi-CAS (3) | New | RIF, INH S |
| *fbiB* | 3642351 | Gly273Arg (ggc/Cgc) | 0.25 | BGD | X-Type (4,1,1,3) | New | RIF, INH S |
| **fbiC** | 1303241 | Tyr104Cys (tat/tGt) | 0.5 | AZE | Beijing (2,2,1) | Retreatment | RIF, INH S |
| **fbiC** | 1303265 | Gly112Ala (ggc/gCc) | 0.12 | BGD | EAI (1,1,3) | New | RIF, INH S |
| **fbiC** | 1303612 | Leu228Phe (ctc/Ttc) | 0.12 | BGD | EAI (1,1,3) | New | RIF, INH S |
| **fbiC-fbiB** | 1303769<br>3642223 | Ser280Leu (tcg/tTg)<br>Arg230Gln (cgg/cAg) | 0.12 | BGD | EAI (1,1,3) | New | RIF, INH S |
| **fbiC** | 1304498 | Pro523Leu (cct/cTt) | 0.5 | BGD | EAI (1,1,3) | New | RIF, INH S |
| **fbiC** | 1305101 | Asn724Ser (aac/aGc) | 0.25 | BGD | Beijing (2,2,1) | New | RIF, INH S |
| **fbiC** | 1305215 | Ser762Asn (agc/aAc) | 0.12 | BGD | Delhi-CAS (3) | Retreatment | RIF, INH S; FQ-R |
| **fbiC** | 1305434 | Ala835Val (gcg/gTg) | 0.5 | BGD | EAI (1,1,3) | New | INH R |
| **fbiC** | 1305494 | Ala855fs (Del 62 nt) | 0.5 | SA | Haarlem<br>(4,1,2,1) | New | RIF, INH S |
| **fbiC-fbiB** | 1305494<br>3642874 | Ala855fs (Del 62 nt)<br>Leu447Arg (ctg/cGg) | 0.25 | SA | mainly-T (4,8) | Retreatment | MDR |
| **fbiC** | 1305496 | Ala856Pro (gcc/Ccc) | 0.25 | BGD | EAI (1,1,3) | New | RIF, INH S |

Overall, the MIC levels among DLM-resistant isolates ranged from 0.12 µg/ml to ≥4 µg/ml, with the highest MIC values occurring with mutation types causing a truncated Ddn or FbiA protein (frameshift or stop codon mutations). The remaining mutations were associated with increased DLM MIC values between 0.12 and 0.5 µg/ml. Opposite to the high MIC level (≥4 µg/ml) observed for the frameshift at codon 14 in the *ddn* gene, the observed frameshift at the very end of the *fbiC* gene (codon 855) caused a lower increase in MIC level at 0.5 µg/ml (Table 2).

Similar to Rv0678, the mutation structural analysis was performed for the Ddn (PDB 3R5L; 3R5R) and Fgd1 (PDB 3B4Y; 3C8N) proteins (Dataset S2). The highly conserved Ddn protein catalyzes the reduction of nitroimidazoles of DLM prodrug by the co-factor F420-H_2_, resulting in intracellular release of lethal reactive nitrogen species (**36**). For the on-target Ddn protein, mutations Asn91Thr and Pro86Thr localize very close to the cofactor binding site (Fig 4A) and ΔDSX energies resulted from DrugScore analysis indicate that these are the only two mutations which reduce binding affinity between Ddn and the cofactor F420-H_2_ (Dataset S2). The Ddn mutation Thr140Ile is far from the cofactor binding site but its effect on MIC increase is due to the protein folding stability, because the side chain of Thr140 residue is involved in the hydrogen bond network with Ala82-Lys79-F420-H_2_ (Fig. 4A). The mutation Val61Gly, which has a mild effect on the MIC (0.12 µg/ml), showed high levels of ΔΔG Kcal/mol with both the Eris and MAESTRO analyses, suggesting a role in destabilizing the folding protein in the β-sheet (Fig. 4A). The analysis of point mutations in Ddn without available MIC values revealed a high level of ΔΔG Kcal/mol energy for mutations Arg72Gln, Pro86Thr and Glu150Ala, suggesting their potential involvement on Ddn stability and consequently phenotypic DLM resistance (Dataset S2).

**Fig. 4.**
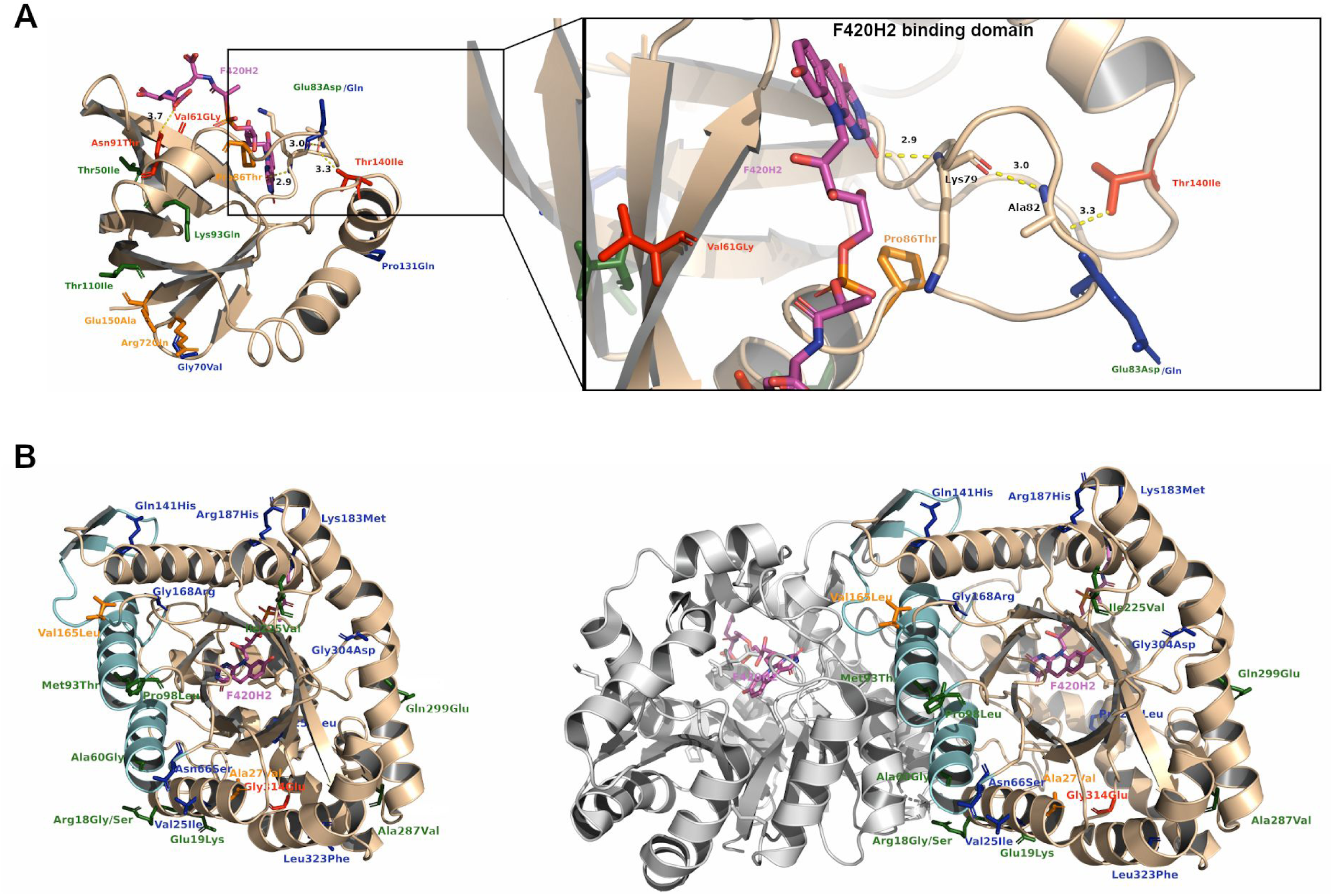
Cartoon representation Ddn and Fgd1 protein structures with mutations found associated to DLM-resistant or associated to DLM-susceptible phenotype. **A**: Ddn protein bound to F420 cofactor (3R5R) with highlighted in sticks the resistant (red), susceptible (green), *in silico* DLM-resistant associated mutations (orange) and *in silico* predicted DLM-susceptible mutations (blue). F420 is shown in magenta licorice. A zoom cartoon representation of Ddn bound to F420 is reported. The hydrogen bond network Thr140-Ala82-Lys79-F420-H2 is shown in dashed yellow line and the Ala82 and Lys79 residues are shown in sticks. **B**: Fgd1 carton representation of holo- and homodimer-Fgd1 bound to F420 (3B4Y). Helixes involved in protein dimerization are colored in cyan. Highlighted in sticks the DLM-resistant (red), DLM-susceptible (green), *in silico* predicted DLM-resistant mutations (orange) and *in silico* predicted DLM-susceptible mutations (blue).

The DLM off-target F420-dependent glucose-6-phosphate dehydrogenase (Fgd1) is important in MTBc energy metabolism, and it is implicated in DLM redox processes related to non-replicating persistence by providing the reduced co-factor F420-H_2_ (**35**). The reported MIC values did not show a strong effect in the *in vitro* experiments because all identified mutations are further than 10 Å from the co-factor F420 binding site (Fig. 4B). Moreover, the computational Eris approach predicts that two Fgd1 mutations without phenotypic data, Ala27Gly and Val165Leu, have potential roles in protein destabilization (Dataset S2). The Fgd1 mutation Gly314Glu which was observed in a strain with only a moderate increase of DLM MIC level seemed to poorly correlate with DLM phenotype, suggesting that other factors could contribute to this small variation of the DLM MIC level in MTBc strains.

The distribution analysis of mutations across genotypes and country of isolation showed that *ddn* mutations involved in a DLM-resistant phenotype are represented only once or twice. Exception are, Pro2Gln which was found in seven mainly-T (4,8) isolates (all from Azerbaijan), Asn91Thr in *ddn* in combination with mutation Val58Ile in *fbiA* in three mainly-T (4,8) isolates (two from Azerbaijan and one from Ukraine), and the high-level resistant stop codon mutation Gln58STOP which was detected in three Beijing (2,2,1) isolates from Ukraine (Fig. 5A). Two DLM-resistant mutations in the *fbiA-fbiB* region were seen in a single isolate, while the *fbiB* mutation Asp224Asn was found in two Delhi-CAS (3) isolates from Bangladesh. Mutation type Ile208Val in *fbiA* was the most prevalent and seen in 22 isolates, all belonging to the Euro-American lineage (Clade1; 4,1,2) and isolated in South Africa (*n=* 19), Bangladesh (*n=* 2) and Ukraine (*n=* 1) (Fig. 5B). Four of the seven DLM-resistant *fbiC* mutation types were seen in single isolates, one was observed in two isolates while two were more prevalent: the frameshift mutation Ala855fs (a deletion of 62 nt) which was detected in 30 isolates from South Africa belonging to eight different lineages, and mutation Ala835Val detected in 18 EAI (1,1,3) isolates from Bangladesh only (Fig. 6). Of note, most of the DLM-resistant strains with high-level of DLM MIC were isolated in Bangladesh (75%) and mutations were detected in *ddn* (46%), *fbiC* (31%), *fbiA* (15%) and *fbiB* (8%) (Dataset S1).

**Fig. 5.**
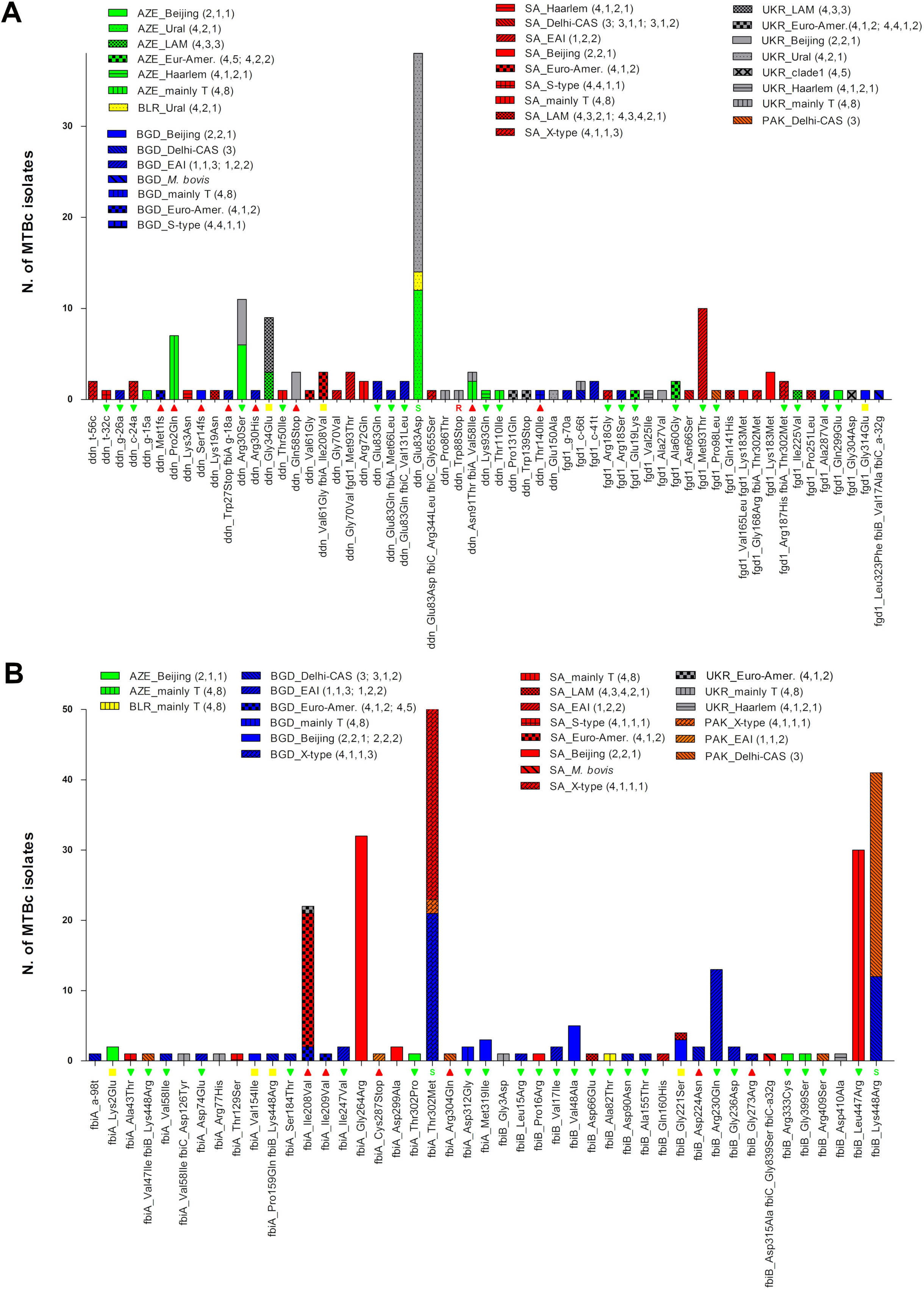
Lineage and country distributions of MTBc strains with variants in *ddn* and *fgd1* genomic regions and *fbiA-fbiB* operon region. The graph reports all mutations found in *ddn, fgd1* **(A)** and *fbiA, fbiB* **(B)** genomic regions, showing their distribution among lineages and country of isolation. The histograms refer to the number of strains in which mutations were observed (Y axis). The colors of the histograms represent the different countries of isolation while the patterns inside each bar represent the different lineages. On the X axis, the results of the MIC test for available MTBc strains are also reported: red triangle are DLM-resistant strains (MIC ≥ 0.12 µg/ml); yellow box are low resistance level (MIC = 0.12 µg/ml) and green triangles are DLM-susceptible strains (MIC < 0.12 µg/ml). If a mutation was previously described in literature it was also reported (“S” for strain susceptible to DLM or “R” for strain resistant to DLM).

**Fig. 6.**
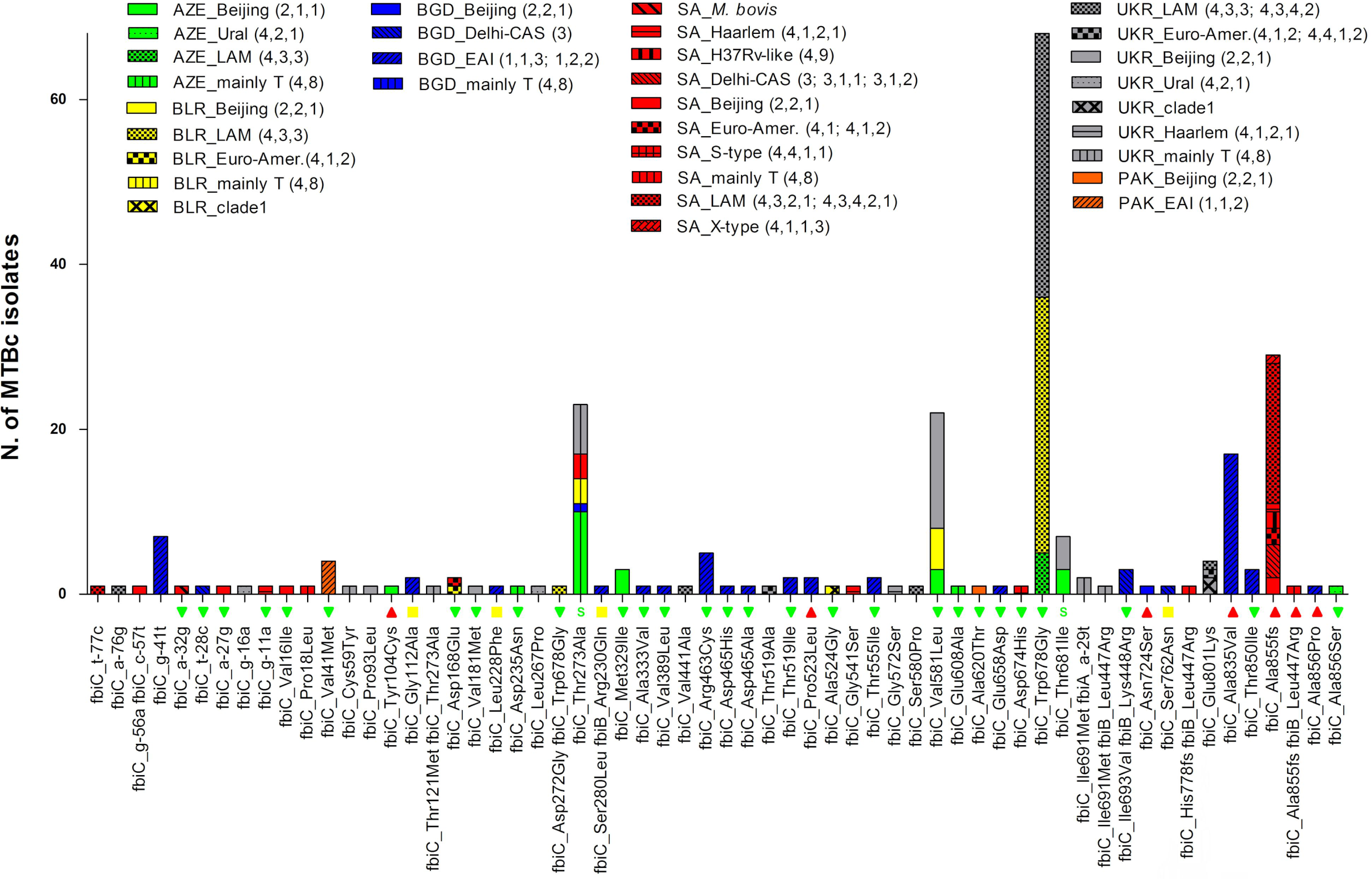
Lineage and country distributions of MTBc strains with variants in *fbiC* genomic region. In this graph are report all mutations found in *fbiC* genomic region showing their distribution among lineages and countries of isolation. See Fig. 5 legend for details.

Cluster analysis by distance matrix was also performed in order to understand if the observation of three or more isolates with mutations associated with DLM resistance were potential clonal clusters. Results showed that the three Beijing (2,2,1) isolates harbouring the stop codon mutation Gln58STOP in *ddn* gene were part of the same transmission chain, at 12 SNPs cut-off (Fig. S2). Moreover, ten other clusters of two isolates each were identified in all the other groups harbouring mutation associated with DLM resistance: four clusters of isolates with the Ile208Val mutation in *fbiA*, one cluster of isolates with Ala855fs variant in *fbiC*, two clusters of isolates with Pro2Gln in *ddn*, one cluster of isolates with Asn91Thr and Val58Ile mutations in *ddn* and *fbiA* respectively, and two clusters of isolates with *fbiC* Ala835Val (Fig. S2).

## 4. Discussion and Conclusion

Bedaquiline (BDQ) and delamanid (DLM) have expanded available treatment options and improved treatment success rates for patients with pulmonary MDR-TB and XDR-TB (**44, 45, 46, 47**), including children with MDR-TB (**48, 49**). The detection of resistance to BDQ and DLM is critical to ensuring effective treatment and care for DR-TB patients and preventing ongoing transmission. Although evidence for the validation and standardization of efficient methods for MICs and the setting of breakpoints for BDQ and DLM continues to expand (**22, 34, 50, 51**), there is still a notable lack of suitable data on resistance-related genomic variants (**52**). Moreover, phenotypic methods are too slow to provide early indication of susceptibility status at the time of treatment initiation. An accurate classification of SNPs according to their association with drug resistance is therefore essential to allow the use of WGS to guide the composition of treatment regimens (**7, 53**). The fact that accurate databases with catalogued mutations are currently lacking for these drugs, represent a serious limitation for molecular DST for BDQ and DLM.

To tackle this, we used a unique dataset containing 4795 WGS data of MTBc isolates from different countries with either a higher burden of TB or MDR-TB (Azerbaijan, Bangladesh, Belarus, South Africa, Pakistan and Ukraine) as a unique and accurate source of genetic information for the characterization and validation of genomic variants potentially involved in BDQ and DLM resistance. In particular, this study highlighted the role of genetic variants for BDQ and DLM resistance development by combining the MICs results of MTBc isolates with variants and the *in silico* analysis on available protein structures, paving the way for the construction of an encyclopaedia of characterized mutations to be use for molecular DST.

From the whole WGS dataset, we identified 61 different BDQ-related variants in *Rv0678, atpE* promoter and *pepQ* genomic regions, out of which four were associated to BDQ-resistant phenotype: two frameshift mutations in Rv0678 associated to a high levels of BDQ MIC and two non-synonymous mutations found associated to low levels of BDQ resistance. To the best of our knowledge, among these four BDQ-resistant mutations, only the frameshift mutation Gly6fs (del_16-17 gg) has been previously described in one BDQ-resistant isolate (**54**). In agreement with the *in vitro* MIC experiments, the *in silico* structural analysis on the Rv0678 single monomer protein form (PDB 4N5B) showed that mutations Met111Lys and Arg96Trp have very high Eris and MAESTRO ΔΔG Kcal/mol energy values, indicating a strong destabilization of the protein folding. The mutations Gly87Arg and Leu117Arg in *Rv0678* were previously described in BDQ-susceptible strains (**13**), confirming the detected low MIC of 0.03 µg/ml and 0.12 µg/ml, respectively. (Dataset S1). The role of Leu117Arg remains unclear as another study described this mutation as associated with both BDQ and also CLF resistance (**11**). However, our phenotypic and *in silico* results suggest that Leu117Arg affects the dimerization of Rv0678 causing a small increase but not high-level value of BDQ MIC. Other mutations in *Rv0678* were described at the same codon position but with a different amino acid change (**5, 8, 9, 11, 13**). The *Rv0678* mutations Val20Phe, Ala84Glu and Arg90Pro were observed to be linked to increase MICs for CLF and potentially associated to a BDQ-resistant phenotype, while Val20Gly was associated with both BDQ and CLF resistance (**5, 9**). In this study, *Rv0678* mutations Val20Ala and Ala84Val were found in BDQ-susceptible strains (MIC ≤ 0.008 µg/ml), while Arg90Cys was associated with a BDQ MIC value of 0.12 µg/ml. Moreover, *in silico* analysis confirmed that the Arg90Cys mutation can have a mild effect on protein stability and a role in the small variation of BDQ MIC. Mutations Arg96Gln, Met146Thr and Leu136Pro *in Rv0678* have been described with increased MIC values for BDQ (**8, 11, 13**). In our dataset, strains harbouring mutations at the same codons were not consistently phenotypically resistant, with mutations Arg96Trp, Met146Arg and Leu136Val respectively showing MICs of 0.25 µg/ml (BDQ-resistant), 0.06 µg/ml (BDQ-susceptible) and 0.03 µg/ml (BDQ-susceptible). Again, *in silico* analysis results were in agreement with the phenotypic data, showing that only the Arg96Trp mutation highly destabilized Rv0678 folding while the other two mutations showed a lower ΔΔG Kcal/mol energy values (Dataset S2). Furthermore, the *in silico* MM/GBSA analysis, revealed that Rv0678 mutations Leu117Arg, Val120Met and Asp141His can affect the protein homodimerization, which could explain the slight increase of BDQ MIC to 0.12 µg/ml for these MTBc strains, yet still classified as BDQ-susceptible despite being close to the proposed cut-off.

Considering the five genomic regions associated with DLM phenotype (*ddn, fgd1, fbiA, fbiB* and *fbiC*), the WGS analysis revealed a total of 164 unique mutations potentially involved in DLM resistance. Apart from seven previously characterized mutations (**16**), all the other 156 DLM-related variants have not been previously described earlier except for Asn91Thr in *ddn*, for which we confirmed its role in DLM resistance (**18**). Phenotypic results on a subset of available isolates showed that 32 different mutations, detected in all of the considered genomic regions were associated to DLM resistance. Notably, the *in silico* mutation structural analysis revealed that the effect of the point mutations in Ddn and Fgd1 were in agreement with the MIC results (Dataset S2). Indeed, the mutation Asn91Thr in Ddn is directly involved in the binding with the co-factor F420-H_2_ by disrupting the Ddn-F420-H_2_ interaction but also in destabilizing the Ddn folding and mutations Val61Gly and Thr140Ile, which were observed in MTBc strains with MICs for DLM of 0.12 µg/ml and 0.5 µg/ml respectively, showed also a potential effect on Ddn protein folding and stability. Furthermore, the *in silico* analysis indicates that three mutations in Ddn for which corresponding phenotypic data were not available (Arg72Gln, Pro86Thr, Glu150Ala) may have a significant impact on protein stability and thereby play a role in the DLM-resistant phenotype.

In addition, the correlation analysis between mutations linked to BDQ- and DLM-resistant phenotypes and metadata information corroborate with previously reported data, suggesting the absence of links between BDQ or DLM resistance and strain lineage or drug resistance profiles of MTBc isolates (**10, 16**). Globally, considering DST profiles of BDQ- and DLM-resistant strains, 75% were fully susceptible, 6% were MDR-TB, and the majority of them (68.7%) was from new TB cases. Moreover, the analysis of country-lineage distribution did not reveal any significant correlation between BDQ/DLM-related mutations and lineage groups or country of isolation. To complete this set of analyses, we also performed a SNP-based distance matrix to evaluate the relatedness of strains harbouring the most frequent DLM-resistant variants, showing that for these groups there are no major transmission chains but only small clusters of two to three isolates, meaning that these resistance-associated mutations can rise spontaneously and independently. Conversely, none of the BDQ-resistant variants were detected in isolates groups.

In conclusion, our study provides novel and important evidence on the role of mutations associated with BDQ- and DLM-resistant phenotypes. A concerningly high prevalence of genetic mutations associated with an increased MIC was detected in clinical isolates from patients who have never been exposed to these drugs, supporting previous findings (**10, 16**). Also, our data showed that different non-synonymous or indel mutation at the same nucleotide position can display a completely different phenotypic effect or different level of resistance, thus reinforcing the need to accurately investigate the role of each individual mutation. Equally important, these findings also showed the presence of 125 genetic variants not associated with BDQ and DLM resistance, scattered over the full length of each target gene. Therefore, considering the complexity of BDQ and DLM mechanisms of resistance and the absence of fully standardized phenotypic tests, the development of accurate molecular-based DST is wholly dependent on the establishment of a complete database of validated mutations, a scenario which is comparable to the challenges associated with molecular markers of resistance to pyrazinamide (PZA) (**55**). The establishment of a common database combining data from MTBc isolates collected in a large number of settings with the inclusion of different parameters (phenotype, genotype, structure, and free energy analyses) is fundamental to improve our understanding of the role of mutations in determining the BDQ/DLM susceptibility phenotype. Finally, this database could be also beneficial to study genetic resistance to other drugs that could be potentially sharing similar genetic basis of resistance such as clofazimine for BDQ and pretomanid for DLM.

For these reasons, despite some limitations (not all strains were available for MIC determination, absence of standardized methods and breakpoints for the interpretation of BDQ and DLM phenotypes, absence of 3D structures of FbiA, FbiB, FbiC and PepQ proteins for *in silico* investigation, lack of knowledge of other genomic regions potentially involved in BDQ/DLM resistance), this work will be an important source of information for new genome-based sequencing approaches for predicting BDQ and DLM resistance.

## Legends

**Table S1.** *M. tuberculosis* genomic regions considered for bedaquiline and delamanid resistances. ^*a*^ For promoter regions, it was considered the upstream region up to the −100 position before the first nucleotide of each gene. ^*b*^ The genomic positions are based on the reference genome of *M. tuberculosis* H37Rv NC_000962.3.

|  | *locus name*<sup>a</sup> | *genomic region*<sup>b</sup> |
| --- | --- | --- |
| **Delamanid (DLM)** | *fbiA_ups* | [3640142-3640542] |
|  | *fbiA* (Rv3261) | [3640543-3641538] |
|  | *fbiB* (Rv3262) | [3641535-3642881] |
|  | *fbiC_ups* | [1302682-1302930] |
|  | *fbiC* (Rv1173) | [1302931-1305501] |
|  | *fgd1_ups* | [490683-490782] |
|  | *fgd1* (Rv0407) | [490783-491793] |
|  | *ddn_ups* | [3986744-3986843] |
|  | *ddn* (Rv3547) | [3986844-3987299] |
| **Bedaquiline (BDQ)** | *atpE_ups* | [1460997-1461044] |
|  | *atpE* (Rv1305) | [1461045-1461290] |
|  | *Rv0678_ups* | [778890-778989] |
|  | *Rv0678* | [778990-779487] |
|  | *pepQ_ups* | [2860419-2860518] |
|  | *pepQ* (Rv2535) | [2859300-2860418] |

**Fig. S1. Study design, number of selected *M. tuberculosis* (MTBc) isolates and DST profile categorization. A**: phenotypic drug susceptibility test (pDST) of the whole collection which classified MTBc isolates in not-MDR, MDR and XDR strains. **B**: Flow chart scheme of isolates selection and stratifications. The blue scheme refers to MTBc isolates selected for DLM-related mutations while the red once to MTBc isolates selected for BDQ-related mutations. ^a^ The WHO approved study list includes 4795 whole genome sequencing (WGS) samples (accession number SRP128089). ^b^ Stratification by the phenotypic resistance profile for the selected isolates. Abbreviations: **not-MDR**, fully susceptible strains (full-S), mono resistant to rifampicin (RIF) or to isoniazid (INH); **MDR**, multidrug resistant strains, resistant almost to INH and RIF; **INH-R**, resistant to isoniazid; **RIF-R**, resistant to rifampicin; **FQ-R**, resistant to fluoroquinolones; **Fully-S**: susceptible to INH and RIF; **Pre-XDR**, MDR resistant also to FQ or second line injectables; **XDR**, MDR plus resistance to FQs and second-line injectables.

**Dataset S1. Samples general database.**

General database with all information of selected MTBc strains harbouring at least one mutation pattern in candidate genes for BDQ and DLM resistance. The excel database is divided in two sheets named “mutations list for BDQ” and “mutations list for DLM”. ^*a*^ WGS analysis (genomic regions for BDQ/DLM resistance) in which are report the list of mutations from WGS analysis with information of genomic locus, gene ID, genomic coordinate (reference strain H37Rv NC_000962.3), amino acid substitution and type of mutation. ^*b*^ MIC value results (µg/mL) from REMA assay (DLM/BDQ) of selected isolates (for several mutations two different isolates were tested); empty cell means that the strain with that specific mutation was not available. ^*c*^ Previously reported mutations: If a mutation was previously reported it is indicated if linked or not to resistance phenotype. ^*e*^ Information on selected samples (isolates 1 and 2): WGS sample name in the WHO database, country of origin, lineage (coll. lineage) and DST profile of MTBc isolates selected for MIC test. ^*d*^ Lineage/Country mutation frequency distribution: numbers of MTBc isolates carrying mutations among countries of origin and strain lineage.

**Dataset S2. Mutation structural analysis.**

General database with all results from the *in silico* analysis of point mutations for the available crystal structures of proteins: Ddn (PDB 3R5R), Fgd1 (PDB 3B4Y) and Rv0678 (PDB 4NB5). Free energy calculation (ΔΔG kcal/mol) of all amino acid change mutations were performed with Eris, an automated estimator of protein stability and MAESTRO, an approach for multi agent stability prediction upon point mutations. DrugScore (DSX) software, a Knowledge-Based Scoring Function for the Assessment of Protein–Ligand Complexes, was used for Ddn-F420 and Fgd1-F420 complexes analysis. The calculation of mutations effect on Rv0678 dimer stability was performed using MMPBSA.py program within Amber14 suite. See materials and methods for details.

**Figure S2. Heatmap of SNP based cluster analysis by distance matrix**. The figure shows the SNP-based cluster analysis results of six MTBc strains groups harbouring the most frequent DLM-resistant related mutations. Colour scale in the square refers to the number of SNPs between each strain (12 SNPs threshold is reported from white to blue). The max number of SNPs are set to 15. MTBc lineages information is also reported.

**Text S1. Supplementary material and methods**. Supplementary information about protocols used for REMA and *in silico* analyses.

